# Susceptibility to auditory feedback manipulations and individual variability

**DOI:** 10.1101/2024.10.02.616332

**Authors:** Muge Ozker, Peter Hagoort

## Abstract

Monitoring auditory feedback from hearing one’s own voice is important for fluent speech production as it enables detection and correction of speech errors. The influence of auditory feedback is best illustrated by manipulating it during speech production. A common temporal manipulation technique, delaying auditory feedback (DAF), leads to disruptions in speech fluency, while a common spectral manipulation technique, perturbing the pitch of auditory feedback (PAF), results in vocal alterations.

Previous research involving clinical populations has revealed diverse susceptibility profiles to auditory feedback manipulations, yet the extent of such diversity within the neurotypical population remains unclear. Furthermore, different types of manipulations elicit distinct speech errors (i.e. fluency/coordination versus acoustic errors), which may be processed by distinct error correction mechanisms. It is yet to be understood whether individuals affected by one manipulation are similarly impacted by the other. Lastly, based on evidence from clinical studies, which demonstrated that visual feedback can improve impaired speech production, it is an open question whether visual feedback can alleviate the disruptive effects of altered auditory feedback.

We recorded voice samples from 40 neurotypical participants during both a DAF and a PAF task. DAF significantly prolonged articulation duration and increased voice pitch and intensity. In some trials, participants received immediate visual feedback, however visual feedback did not alleviate but rather strengthened the disruptive effects of DAF. During the PAF task, participants adjusted their voice pitch in the opposite direction of the perturbation in majority of the trials to compensate for the perturbation. We assessed susceptibility of the participants to the effects of DAF and PAF by examining articulation duration and compensatory vocal response magnitude, respectively. Susceptibility varied widely among participants for both manipulations, but individuals susceptible to one manipulation did not consistently exhibit susceptibility to the other, indicating distinct processing mechanisms for these different types of auditory feedback manipulations.

## INTRODUCTION

Auditory feedback from hearing one’s own voice during speech production plays an important role in speech monitoring as it allows for the detection and instant correction of vocalization errors. Disruption of auditory feedback affects vocalization, as evidenced by how post lingually deafened individuals may develop speech fluency problems (1, 2). The influence of auditory feedback on speech is usually demonstrated by various temporal or acoustic manipulations. Delaying auditory feedback (DAF) during speech production is recognized to cause a decrease in speech rate and induce dysfluencies reminiscent of stuttering (3-5). Likewise, perturbing the fundamental frequency of auditory feedback (PAF) leads to slight changes in the voice pitch, typically in the opposite direction of the perturbation as a compensatory response (6, 7).

Previous studies in clinical populations have reported variable susceptibility profiles to auditory feedback manipulations. Schizophrenic patients with auditory hallucinations (8) and individuals with autism spectrum disorder have demonstrated heightened susceptibility to the effects of DAF (9). Conversely, patients with conduction aphasia exhibit lower susceptibility to DAF (10). While delaying auditory feedback results in stutter-like speech in neurotypical individuals (5), it has been found to improve fluency in individuals who stutter (11). For PAF, compensatory vocal responses to PAF were shown to be markedly reduced in post-stroke aphasia patients (12), whereas they were heightened in Parkinson’s disease patients (13) compared to healthy individuals. In order to understand this wide range of susceptibility in clinical populations, it is essential to initially assess the variability within healthy populations. Several studies involving neurotypical individuals have also documented variable susceptibility to DAF (14-17) and PAF (18, 19), however the extent and the underlying cause of this variability remain unclear.

One issue is that these two most widely used auditory feedback manipulations impact speech in entirely dissimilar manners, eliciting distinct type of speech errors. DAF introduces a temporal discrepancy between articulation and auditory feedback, resulting in coordination errors (timing, rate, fluency, articulation), while PAF alters the spectral components of auditory feedback leading to errors in vocal acoustics. If temporal and spectral manipulations trigger distinct error correction mechanisms, it remains uncertain whether individuals who are affected by one type of manipulation will also be affected by the other. Susceptibility could be specific to a particular aspect of auditory feedback or it could represent a more general phenomenon.

Another question is why certain individuals exhibit heightened susceptibility to auditory feedback manipulations, while others demonstrate the ability to withstand them. One proposed hypothesis suggests that individuals may exhibit varying sensory dominance, indicating a reliance on one sensory modality over another (20). During speech production, individuals receive sensory feedback from multiple sources, including auditory feedback (hearing their own voice), proprioceptive feedback (from muscles and joints involved in articulation), and somatosensory feedback (from oral structures like lips and tongue). Individuals who are more susceptible to the effects of auditory feedback manipulations may lean heavily on auditory feedback, whereas those less susceptible may utilize alternative sensory feedback sources to a greater extent. Evidence from clinical populations demonstrate that immediate visual feedback can improve impaired speech production (21, 22). We predicted that the presence of visual feedback during speech production with manipulated auditory feedback could potentially improve speech fluency, especially in individuals who exhibit lower susceptibility to auditory feedback manipulations.

In this study we addressed these questions by examining the susceptibility to both delayed and pitch-perturbed auditory feedback (DAF and PAF) in a gender balanced, neurotypical population. We evaluated the major speech output measures that were shown to be affected in previous studies, including articulation duration, voice intensity and pitch, and tested whether immediate visual feedback can ameliorate speech fluency in the context of using DAF. In addition, we also tested whether prolonged exposure to manipulated auditory feedback results in task habituation. Finally, we assessed the interindividual variability in susceptibility to auditory feedback manipulations and tested whether susceptibility profiles are similar for temporal and acoustic manipulations.

## MATERIALS and METHODS

### Participants

40 native Dutch speakers (20 females, age: 24.5 ± 5) without a known history of hearing or language impairments were included in the study after providing written informed consent prior to the start of the experiments (Participant recruitment period: 13/04/2022 - 01/08/2022). The research was approved by Radboud University’s Faculty of Social Sciences (ethics application ECSW-2020-046) and complied with the Declaration of Helsinki.

### Delayed Auditory Feedback Experiment

#### Stimuli and Equipment

Audio recordings of 10 sentences spoken by a female native Dutch speaker were presented to the participants. The sentences consisted of 10-12 words and had a mean duration of 4 seconds. Stimulus presentation was controlled by Matlab Psychtoolbox-3 running on a Windows computer. Participants’ voice was recorded using the laptop’s microphone, delayed using Matlab Audiotoolbox function and played back to the participants through Sennheiser headphones. Webcam was used to present the video signal of participants’ face. Participants’ voice was also recorded by a Zoom H6 recorder for high quality recordings.

#### Task

Participants were instructed to listen to the auditorily presented sentences and repeat them out loud. Auditory feedback was presented either simultaneously or with 200 ms delay (DAF). In half of the trials visual feedback (VF) was presented in addition to the auditory feedback. Thus, the experiment consisted of four different conditions: no DAF, DAF, no DAF + VF and DAF + VF. For trials with no visual feedback, participants fixated on a crosshair presented in the middle of the screen. For trials with visual feedback, participants were instructed to look at their mouth. Each sentence was repeated multiple times, twice for each condition, which resulted in a total of 80 trials, presented randomly in two consecutive experiment blocks.

#### Analysis

Participants’ audio recordings were analyzed in Praat. For each trial, sentence onset and offset were manually marked to calculate articulation duration. Voice intensity and voice pitch contours were calculated using automated scripts.

#### LME Model

Linear mixed effects models (lmer function in R) were used to test the effects of four experimental manipulations (auditory feedback delay: 0 or 200 ms, visual feedback: present or absent, gender and trial order) on three continuous variables (articulation duration, voice intensity and voice pitch). Because different individuals may have different baseline speech rates or vocal characteristics, or may be affected differently by experimental manipulations, participants were added as a random effect using a random intercept and slope model. In addition, stimulus exemplars were also added as a random effect using a random intercept and slope model to account for any different effects that different sentence stimuli may have on the dependent variables. For ease of interpretation, three fixed effects (delay, visual feedback and gender) were centered around zero, and one fixed effect (trial order) was scaled before model estimation. Models were estimated using REML, and the degrees of freedom (df), t-values, and p-values were calculated according to the Satterthwaite approximation (23).

### Perturbed Auditory Feedback Experiment

#### Stimuli and Equipment

Auditory feedback was perturbed during phonation by shifting the voice fundamental frequency (F0: pitch) by either 100 or 200 cents in the upward or downward direction for 300 milliseconds, starting at a random latency (1400 to 1900 milliseconds) following the phonation onset. Stimulus presentation was controlled by Matlab Psychtoolbox-3 running on a Windows computer. Participants’ voice was recorded using the laptop’s microphone, frequency perturbation was applied using the Audapter software (Cai, Boucek, Ghosh, Guenther, & Perkell, 2008) and auditory feedback was played back to the participants through Sennheiser headphones.

#### Task

Participants phonated the vowel /a/ for 3.5-4 seconds. A text “Aaah” was presented on the screen to prompt the participants to phonate. Auditory feedback was presented back to the participant either with or without perturbation. In trials with perturbation, auditory feedback was perturbed by either 100 or 200 cents in the upward or downward direction. Thus, the experiment consisted of five different conditions: no PAF, +100, +200, -100 and -200 PAF. 30 trials were recorded for each condition, resulting in a total of 150 trials.

#### Analysis

Pitch of the participants’ voice were determined using the autocorrelation method in Praat (24). Pitch contours for each trial were exported to Matlab for further analysis. Trials that contained sharp discontinuities in their pitch track due to algorithm failure were excluded from the analysis. Pitch contours were epoched from -200 to +800 milliseconds with respect to the perturbation onset and were converted from Hertz to cents scale using the following formula:

F0(cents) = 1200 x log_2_ (F / F_baseline_)

Average pitch from -100 to 0 milliseconds before the perturbation onset was used as the baseline (F_baseline_). Total vocal response magnitudes were calculated by averaging the pitch contours for each trial within the +200 to +500 milliseconds time window after perturbation. Each perturbation trial was first classified as having an opposing or a following response by performing linear regression on the pitch contour from +50 to +350 milliseconds after perturbation. Response direction was determined by the slope of the fitted line, with positive slope indicating a following and negative slope indicating an opposing response for the upward perturbation condition and vice versa for the downward perturbation condition as in (25).

#### LME Model

Linear mixed effects models were used to test the effects of four experimental manipulations (perturbation direction: up or down, perturbation magnitude: 100 or 200 cents, gender and trial order on a continuous variable (vocal response magnitude) and a discrete variable (vocal response direction: following or opposing). Vocal response magnitude for each trial was calculated by averaging the rectified pitch contour within the +200 to +500 milliseconds time window after perturbation. A linear mixed effects model (lmer in R) was fit to predict vocal response magnitude and a logistic mixed effects model (glmer in R) was fit to predict vocal response direction. Because different individuals may have different baseline vocal characteristics, or may be affected differently by experimental manipulations, participants were added as a random effect using a random intercept and slope model. To calculate each participant’s susceptibility to perturbed auditory feedback (*see Susceptibility Index Calculation*) an LME model was fit to predict vocal response magnitude using perturbation (present or absent regardless of perturbation magnitude and direction), gender and trial order as fixed effects and participants as a random effect. For ease of interpretation, three fixed effects (perturbation direction, perturbation magnitude and gender) were centered around zero, and one fixed effect (trial order) was scaled before model estimation. Linear mixed models were estimated using REML, and the degrees of freedom (df), t-values, and p-values were calculated according to the Satterthwaite approximation. Generalized linear mixed model was fit by maximum likelihood estimation using Laplace approximation. Z-values and p-values were computed using Wald z-distribution approximation.

### Susceptibility Index Calculation

Linear mixed effect models were fit to articulation duration and vocal response magnitudes for the delayed and perturbed auditory feedback experiments, respectively (*see LME Model*). In these models, participants were added as random factors using a random intercept and slope. Individual susceptibilities to delayed and perturbed auditory feedback were measured by extracting the coefficients of the fitted models and using the slope values for each participant as susceptibility indices for the two manipulations.

## RESULTS

### Delayed auditory feedback experiment

In the delayed auditory feedback experiment, participants repeated out loud auditorily presented sentences. Their voice recordings were analyzed to extract three vocal characteristics including articulation duration, voice intensity and voice pitch (**Figure 1**).

**Figure 1:**
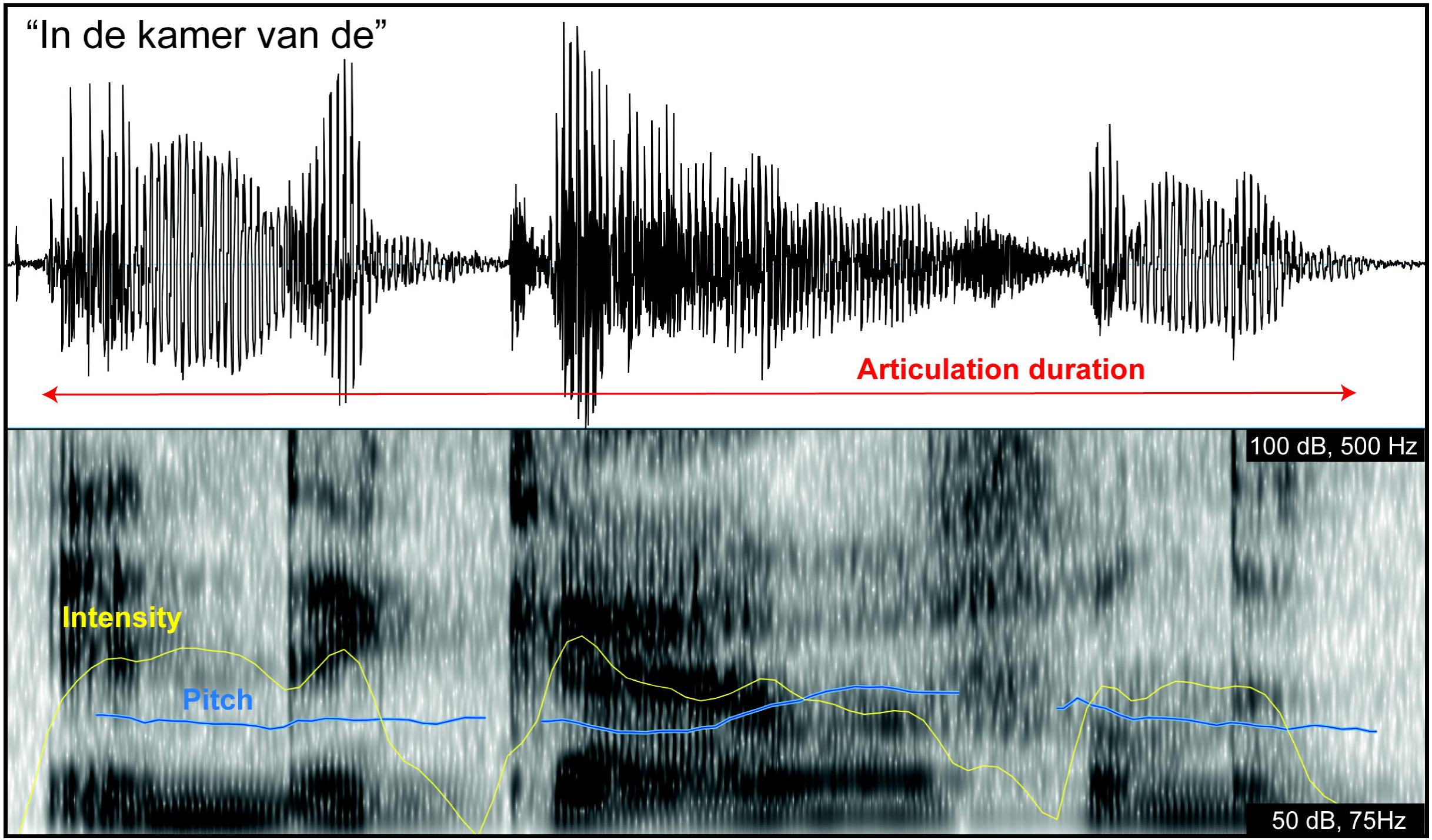
Voice recording analysis. Audio signal (top panel) recorded during the articulation of a sentence. Spectrogram of the audio signal (bottom panel) with darker bands representing the fundamental frequency (pitch) and formant frequencies of the voice. Voice intensity (yellow line) and pitch contour (blue line) are extracted in Praat.

Linear mixed effects models were fit to these vocal response variables to examine the effects of auditory feedback delay, visual feedback, gender and trial order. To account for overall differences across participants and stimulus exemplars, these were added to the models as random factors (*see Methods*). There was a significant positive effect of delay on all vocal characteristics, suggesting that hearing one’s own voice with 200ms delay increases articulation duration (No DAF: 3.2± 0.5 *vs.* DAF: 4.1 ± 1.5 seconds), voice intensity (No DAF: 58.2 ± 7.8 *vs.* DAF: 59.7 ± 7.7 decibels) and voice pitch (No DAF: 156.6 ± 41.3 *vs.* DAF: 159.1± 41.6 Hertz) (**Figure 2a-c**, **Table 1-3**). Also, there was a significant positive effect of visual feedback on all vocal characteristics, suggesting that seeing one’s own face during speech production increases articulation duration (No VF: 3.6 ± 1.1 *vs.* VF: 3.7 ± 1.2 seconds), voice intensity (No VF: 58.8 ± 7.7 *vs.* VF: 59.1 ± 7.8 decibels) and voice pitch (No VF: 157.5 ± 41.5 *vs.* VF: 158.1 ± 41.4 Hertz), reinforcing the disruptive effects of DAF on speech. The interaction of delay and visual feedback was not significant, suggesting that visual feedback affected these voice characteristics regardless of auditory feedback delay.

**Figure 2:**
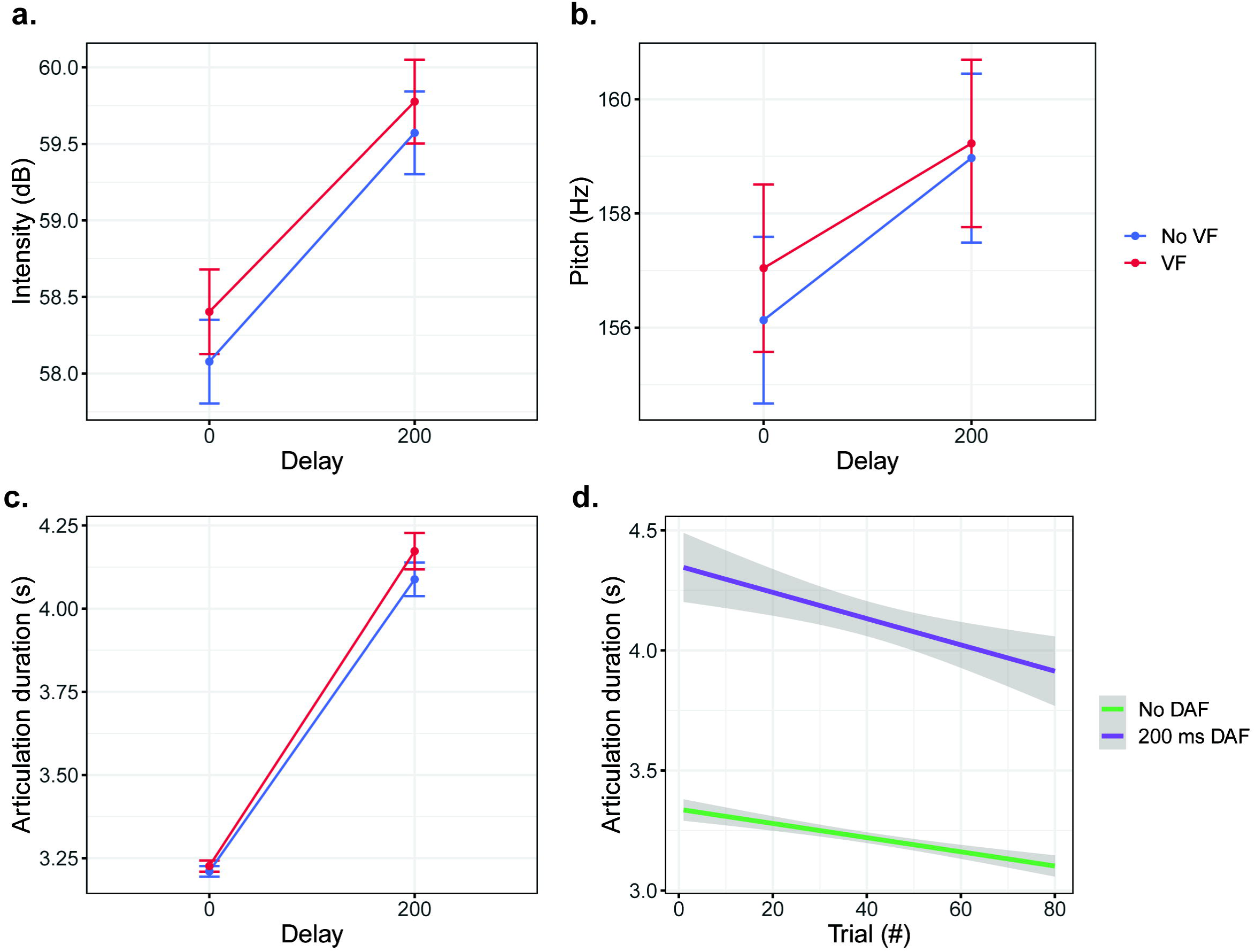
Effect of delayed auditory feedback on vocal characteristics. **a)** Voice intensity, **b)** Voice pitch and **c)** Articulation duration averaged across trials and participants for simultaneous (0ms) and delayed (200ms) auditory feedback when visual feedback is present (red) and absent (blue). Error bars indicate standard error of the mean across trials. d) The effect of trial order on articulation duration for simultaneous (green) and 200ms delayed (purple) auditory feedback. Shaded regions indicate 95% confidence intervals.

**Table 1:** Results of a linear mixed effects model of the articulation duration.

| *Fixed Effects* | *Estimate* | *SE* | *df* | *t* | *p* |
| --- | --- | --- | --- | --- | --- |
| (Intercept) | 3.67 | 0.15 | 43.86 | 24.85 | < 2e-16 |
| **Delay** | **0.91** | **0.19** | **39.58** | **4.67** | **3.4x10<sup>-5</sup>** |
| **VF** | **0.05** | **0.02** | **39.07** | **2.32** | **0.02** |
| Gender | 0.28 | 0.25 | 38.00 | 1.14 | 0.26 |
| **Trial order** | **-0.08** | **0.02** | **32.70** | **-4.93** | **2.3 x10<sup>-5</sup>** |
| Delay x VF | 0.06 | 0.03 | 38.94 | 1.88 | 0.07 |

The fixed effects were delay (0 vs 200 ms), visual feedback (presence or absence), gender, trial order and interaction of delay and visual feedback. Participants and stimulus exemplars were included in the model as random factors. For each effect, the model estimates (in units of seconds) for that factor are shown relative to intercept. Significant effects are shown in **bold**.

**Table 2:** Results of a linear mixed effects model of the voice intensity.

| *Fixed Effects* | *Estimate* | *SE* | *df* | *t* | *p* |
| --- | --- | --- | --- | --- | --- |
| (Intercept) | 58.96 | 1.00 | 41.60 | 58.88 | < 2e-16 |
| **Delay** | **1.41** | **0.21** | **41.42** | **6.71** | **3.9x10<sup>-5</sup>** |
| **VF** | **0.26** | **0.06** | **39.04** | **3.92** | **0.0003** |
| **Gender** | **8.89** | **1.95** | **38.00** | **4.55** | **5.3x10<sup>-5</sup>** |
| Trial order | 0.09 | 0.09 | 39.14 | 1.01 | 0.32 |
| Delay x VF | -0.12 | 0.11 | 3008.4 | -1.05 | 0.29 |

The fixed effects were delay (0 vs 200 ms), visual feedback (presence or absence), gender, trial order and interaction of delay and visual feedback. Participants and stimulus exemplars were included in the model as random factors. For each effect, the model estimates (in units of decibels) for that factor are shown relative to intercept. Significant effects are shown in **bold**.

**Table 3:** Results of a linear mixed effects model of the voice pitch.

| *Fixed Effects* | *Estimate* | *SE* | *df* | *t* | *p* |
| --- | --- | --- | --- | --- | --- |
| (Intercept) | 157.86 | 2.75 | 43.45 | 57.39 | < 2e-16 |
| **Delay** | **2.42** | **0.54** | **36.16** | **4.44** | **8.2x10<sup>-5</sup>** |
| **VF** | **0.54** | **0.23** | **38.68** | **2.32** | **0.026** |
| **Gender** | **75.04** | **5.28** | **38.01** | **14.22** | **< 2e-16** |
| Trial order | -0.48 | 0.33 | 37.02 | -1.48 | 0.15 |
| Delay x VF | -0.61 | 0.50 | 8.99 | -1.22 | 0.25 |

The fixed effects were delay (0 vs 200 ms), visual feedback (presence or absence), gender, trial order and interaction of delay and visual feedback. Participants and stimulus exemplars were included in the model as random factors. For each effect, the model estimates (in units of Hertz) for that factor are shown relative to intercept. Significant effects are shown in **bold**.

Gender did not have a significant effect on articulation duration but it had a significant effect on voice intensity (Male: 54.5 ± 7.5 *vs.* Female: 63.4 ± 4.9 decibels) and pitch (Male: 120.3 ± 15.9 *vs.* Female: 195.4 ± 19.3 cents) as expected, since females have a higher voice pitch and intensity compared to males. Trial order had a significant negative effect on articulation duration but no effect on voice pitch and intensity, suggesting that participants spoke faster as the experiment progressed, but they maintained a stable vocal characteristic throughout the experiment (**Figure 2d**).

Common effects of DAF include increased voice intensity and pitch, which were previously attributed to heightened muscular tension as individuals seek to counteract the experimental interference (4). Additionally, during DAF, individuals often slow down or reset their articulation in an effort to synchronize their speech with the delayed feedback, resulting in prolonged articulation duration (4, 5). As these latter actions are indicative of compensatory attempts to correct speech errors, we focused on articulation duration, rather than voice intensity and pitch, to evaluate the susceptibility profiles of different individuals to DAF. As an example, **Figure 3a** shows the articulation duration from two participants during sentence production with simultaneous and delayed auditory feedback. While delayed feedback caused only a small increase in articulation duration in one participant (Participant #7; No DAF: 3.04 ± 0.27 *vs.* DAF: 3.2 ± 0.32), it caused a much larger increase in another participant (Participant #22; No DAF: 3.04 ± 0.5 *vs.* DAF: 4.9 ± 0.94). To describe this interindividual variability in susceptibility to delayed auditory feedback across participants, the LME model of articulation duration was used. For each participant, slope value of the fixed effect ‘auditory feedback delay’ was used as a susceptibility index. Across participants, there was a considerable variability in susceptibility to delay, measured by the increase in articulation duration during 200ms DAF compared to no DAF condition (slope values ranging 0.14 to 6) (**Figure 3b**).

**Figure 3:**
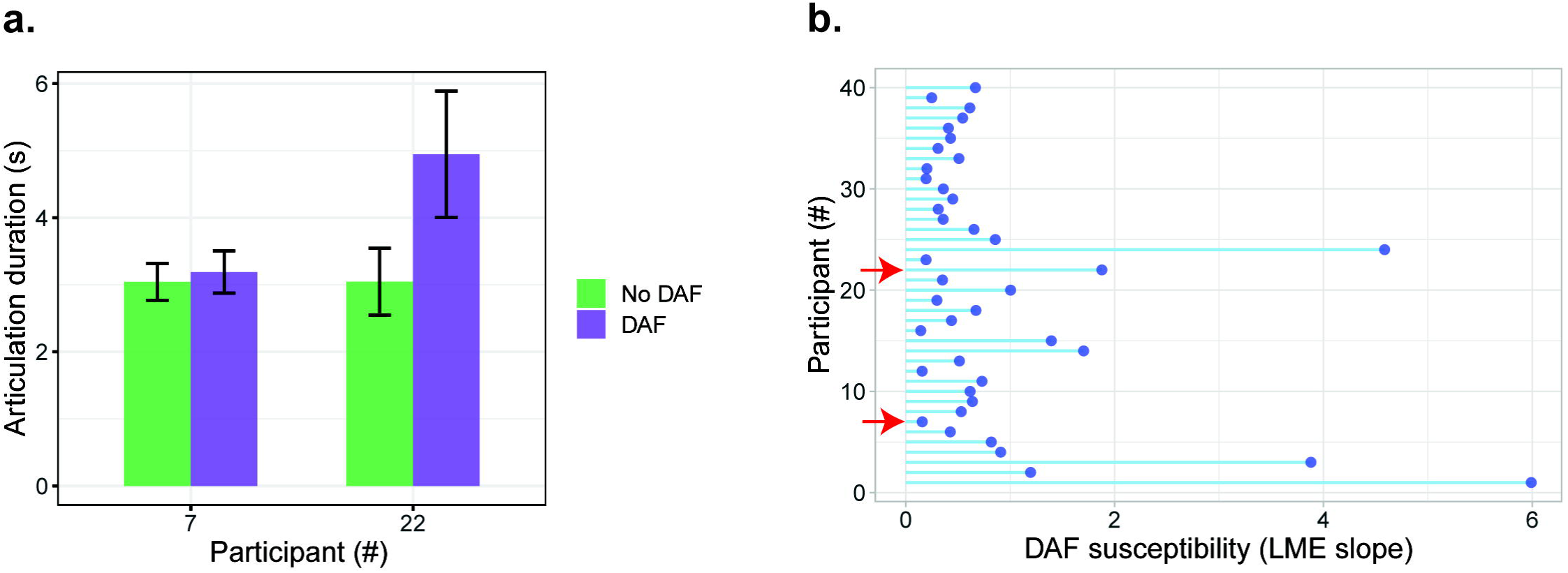
Individual variability in susceptibility to delayed auditory feedback. **a)** Comparison of articulation durations during simultaneous (green bars) and 200ms delayed (purple bars) auditory feedback in two different participants. **b)** Slopes values from the LME model which represent the effect of auditory feedback delay on articulation duration for different participants. Red arrows indicate the participants that were shown in panel a.

### Perturbed auditory feedback experiment

In the perturbed auditory feedback experiment, participants phonated the vowel /a/ for 3.5-4 seconds. During phonation, the fundamental frequency (pitch) of their voice was perturbed 200 cents either in the upward or the downward direction for 300ms. The voice recordings were analyzed to understand how participants responded to perturbed auditory feedback. Examining the average pitch contours revealed that participants slightly shifted their voice pitch in the opposite direction of the perturbation (**Figure 4**). These compensatory vocal responses reached a maximum at ∼520 milliseconds following the perturbation onset, and they were very small compared to the perturbation magnitudes (Mean across participants: 7.7 cents (-100), 8.6 cents (+100), 5.6 cents (-200) and 10.5 cents (+200)).

**Figure 4:**
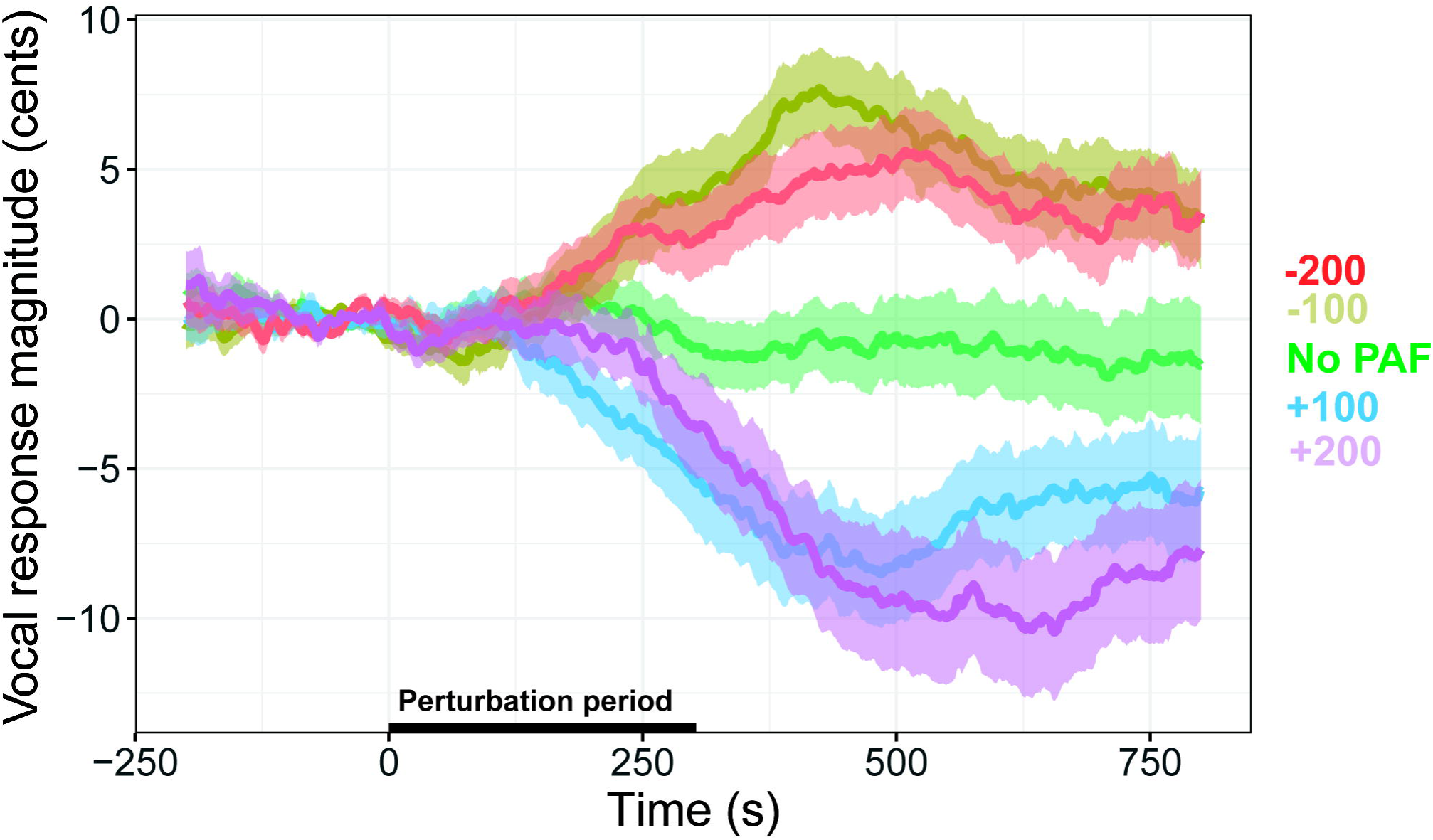
Compensatory vocal responses to perturbation. Pitch contours during the phonation of the vowel /a/ with no perturbation, 100-cent and 200-cent perturbation in the upward and downward directions averaged across participants. Shaded regions indicate standard error of the mean across participants.

Even though the average vocal responses were in the opposite direction of the perturbation, closely inspecting individual trials revealed that participants not always opposed but also followed the perturbation by shifting their voice pitch in the same direction of the perturbation. Categorizing each trial as opposing or following (*see Methods*) confirmed that while the majority of vocal responses were opposing, still a substantial percentage were following responses (Mean across participants: 37.4% (-100), 39.2% (+100), 45.9% (-200) and 43.7% (+200)).

Averaging pitch contours of opposing and following responses separately revealed a fluctuating pattern of vocal responses. For the opposing responses (**Figure 5a**), there were different response phases after the perturbation onset. In the first phase, immediately after perturbation onset (**T1**), responses slightly increased in the same direction with the perturbation. However, in the second phase (**T2**), responses changed direction and increased in the opposite direction of the perturbation. The peak vocal response magnitude was observed in this second phase. In the last phase (**T3**), response magnitudes slightly decreased and remained sustained.

**Figure 5:**
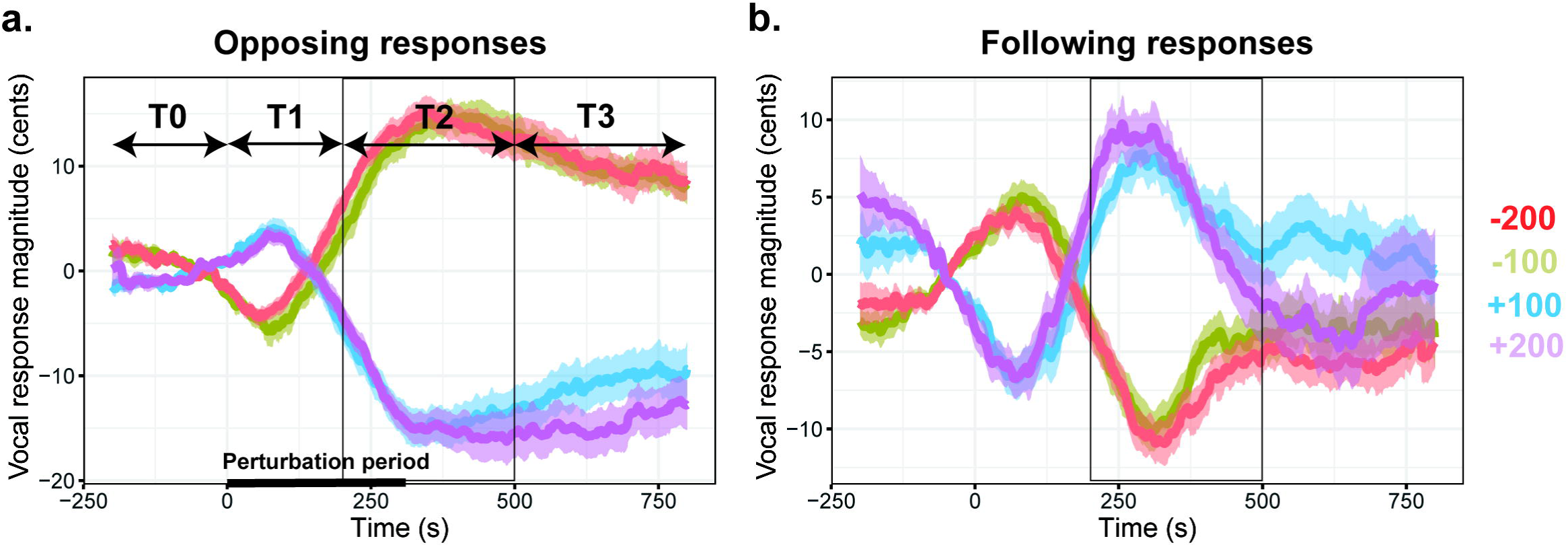
Vocal responses that oppose and follow the direction of perturbation. **a)** Pitch contours of compensatory (opposing) vocal responses during the phonation of the vowel /a/ with 100-cent and 200-cent perturbation in the upward and downward directions averaged across participants. Shaded regions indicate standard error of the mean across participants. T0: - 200 to 0, T1: 0:200, T2: 200:500 and T3: 500 to 800 milliseconds indicate different time periods with respect to the perturbation onset. Black rectangle highlights time period T2 within which the vocal responses are averaged for the LME analysis **b)** Pitch contours of following vocal responses averaged across participants with shaded regions indicating standard error of the mean.

For the following responses (**Figure 5b**), there were also different response phases after the perturbation onset but the reverse pattern was observed. In the first phase, responses slightly increased in the opposite direction with the perturbation. Then, in the second phase, responses changed direction and increased in the following direction of the perturbation. The peak vocal response magnitude was again observed in this second phase. In the last phase, responses remained stable after approaching the baseline levels. Interestingly, there was a significant difference in responses to downward versus upward perturbation directions even before perturbation started (**T0**) for both opposing (t=3.5, p=0.0005, unpaired Welch two-sample t-test) and following responses (t=4.99, p=6.7x10^-7^).

Since opposing responses reflect vocal compensation and error correction, the remainder of the analyses focused on opposing responses. A linear mixed effects model was fit to the vocal response magnitudes averaged in the 200 to 500 milliseconds time window (**T2 in Figure 5a**) to examine the effects of perturbation direction (up or down), perturbation magnitude (100 or 200 cents), gender and trial. To account for overall differences across participants, participant was added to the model as a random factor. None of the effects were significant (**Table 4**), however there was a marginal negative effect of trial order on the response magnitude (p=0.05), which suggests that participants made smaller compensatory vocal responses for the perturbation after repeated exposure to perturbed auditory feedback (**Figure 6b**).

**Figure 6:**
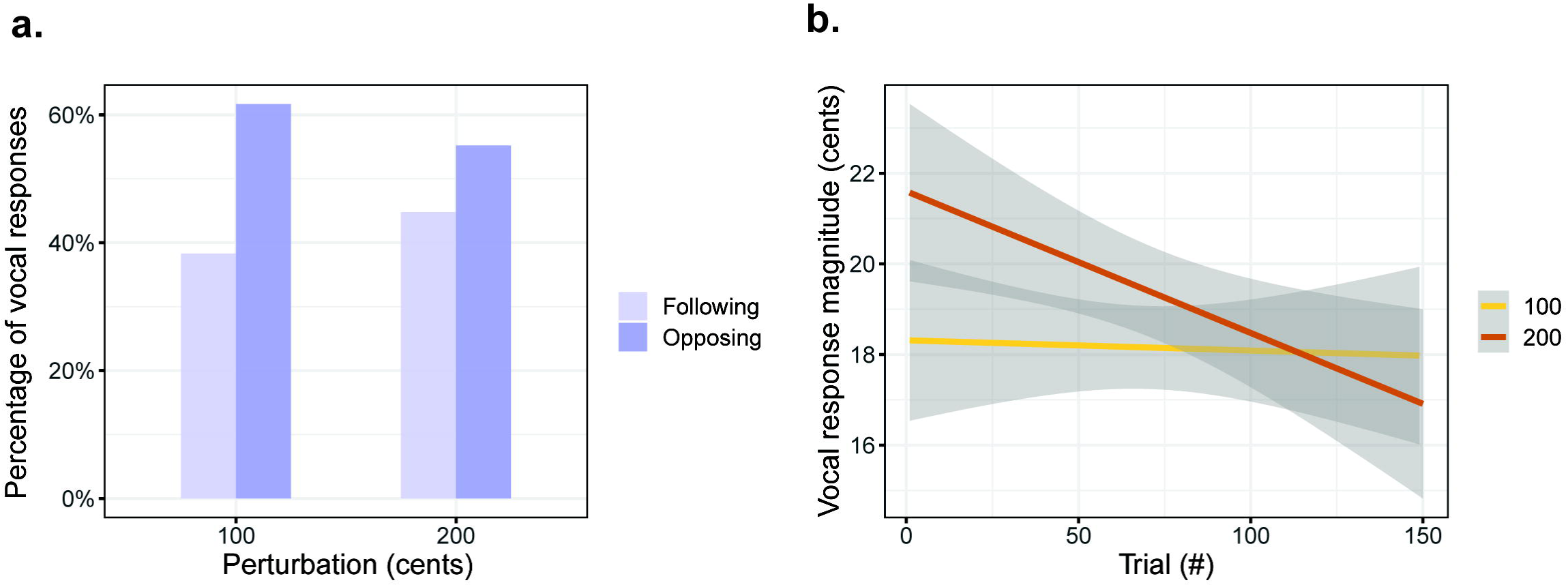
Effects of perturbation magnitude. **a)** Percentage of opposing (dark purple) and following (light purple) vocal responses for 100-cent and 200-cent perturbations. **b)** Change in vocal response magnitudes with the progression of trials for 100-cent (yellow) and 200-cent (brown) perturbation. Shaded regions indicate 95% confidence intervals.

**Table 4:** Results of a linear mixed effects model of the compensatory response magnitude.

| Fixed Effects | Estimate | Std. Error | df | t | p |
| --- | --- | --- | --- | --- | --- |
| (Intercept) | 18.43 | 1.13 | 38.14 | 16.27 | <2e-16 |
| Perturbation magnitude | 0.99 | 0.63 | 2603.4 | 1.56 | 0.12 |
| Perturbation direction | 0.76 | 0.89 | 31.43 | 0.85 | 0.4 |
| Gender | -2.80 | 2.26 | 38.12 | -1.24 | 0.22 |
| Trial order | -0.62 | 0.32 | 2610.3 | -1.95 | 0.051 |

The fixed effects were the perturbation magnitude (100 vs. 200 cents), perturbation direction (upward or downward), gender and trial order. Participants were included in the model as a random factor. For each effect, the model estimates (in units of cents) for that factor are shown relative to intercept.

A general linear mixed effects model was fit to the response direction (opposing or following) to examine the effects of perturbation direction (up or down), perturbation magnitude (100 or 200 cents), gender and trial order. To account for overall differences across participants, participant was added to the model as a random factor. There was a significant effect of perturbation magnitude (z = 4.6, p = 4.6x10^-6^; **Table 5**) driven by a larger percentage of opposing responses for 100-cent compared to 200-cent pitch perturbations in both directions (%62 *vs*. %55; **Figure 6a**). A close inspection of the marginal effect of trial order on response magnitude revealed that this effect was driven by responses to the 200-cent perturbation. Participants reduced their vocal responses mainly for the 200-cent perturbation after repeated exposure to the perturbation (**Figure 6b**).

**Table 5:** Results of a logistic mixed effects model of the vocal response direction.

| *Fixed Effects* | *Estimate* | *Std. Error* | *z* | *p* |
| --- | --- | --- | --- | --- |
| (Intercept) | -0.37 | 0.09 | -4.15 | $3.3 \times 10^{-5}$ |
| **Perturbation magnitude** | **0.29** | **0.06** | **4.58** | **$4.6 \times 10^{-5}$** |
| Perturbation direction | -0.01 | 0.15 | -0.09 | 0.92 |
| Gender | 0.08 | 0.18 | 0.42 | 0.67 |
| Trial order | 0.001 | 0.0007 | 1.76 | 0.08 |

The fixed effects were the perturbation magnitude (100 vs. 200 cents), perturbation direction (upward or downward), gender and trial order. Participants were included in the model as a random factor. For each effect, the model estimates for that factor are shown relative to intercept, with positive estimate values indicating an increase in following responses. Significant effects are shown in **bold**.

Since perturbation direction (up or down) or perturbation magnitude (100 or 200 cents) did not have a significant effect on the vocal response magnitude (**Table 4**), all trials with a perturbation were combined regardless of perturbation magnitude or direction. A linear mixed effects model was fit to the vocal responses to examine the effects of perturbation (non-perturbed or perturbed), gender and trial order. Again, participants were added to the model as a random factor. There was a significant effect of perturbation on the vocal response magnitude (t = 3.27, p = 0.002; **Table 6**).

**Table 6:** Results of a linear mixed effects model of the compensatory response magnitude to all perturbations.

| Fixed Effects | Estimate | Std. Error | df | t | p |
| --- | --- | --- | --- | --- | --- |
| (Intercept) | 17.17 | 0.98 | 38.34 | 17.51 | < 2e-16 |
| **Perturbation** | **2.77** | **0.85** | **38.77** | **3.27** | **0.002** |
| Gender | -3.87 | 1.95 | 37.98 | -1.98 | 0.054 |
| Trial order | -0.33 | 0.25 | 3740.7 | -1.29 | 0.19 |

The fixed effects were the perturbation (presence or absence), gender and trial order. Participants were included in the model as a random factor. For each effect, the model estimates (in units of cents) for that factor are shown relative to intercept. Significant effects are shown in **bold**.

As an example, **Figure 7a** shows the vocal response magnitudes from two participants during phonation with perturbed and non-perturbed auditory feedback. While perturbation did not affect vocal response magnitude in one participant (Participant #5; No Perturbation: 15.8 ± 12.4 *vs.* Perturbation: 16.4 ± 16.8), it caused a significant increase in the other participant (Participant #27; No Perturbation: 16.3 ± 15.4 *vs.* Perturbation: 22.7 ± 17.4).

**Figure 7:**
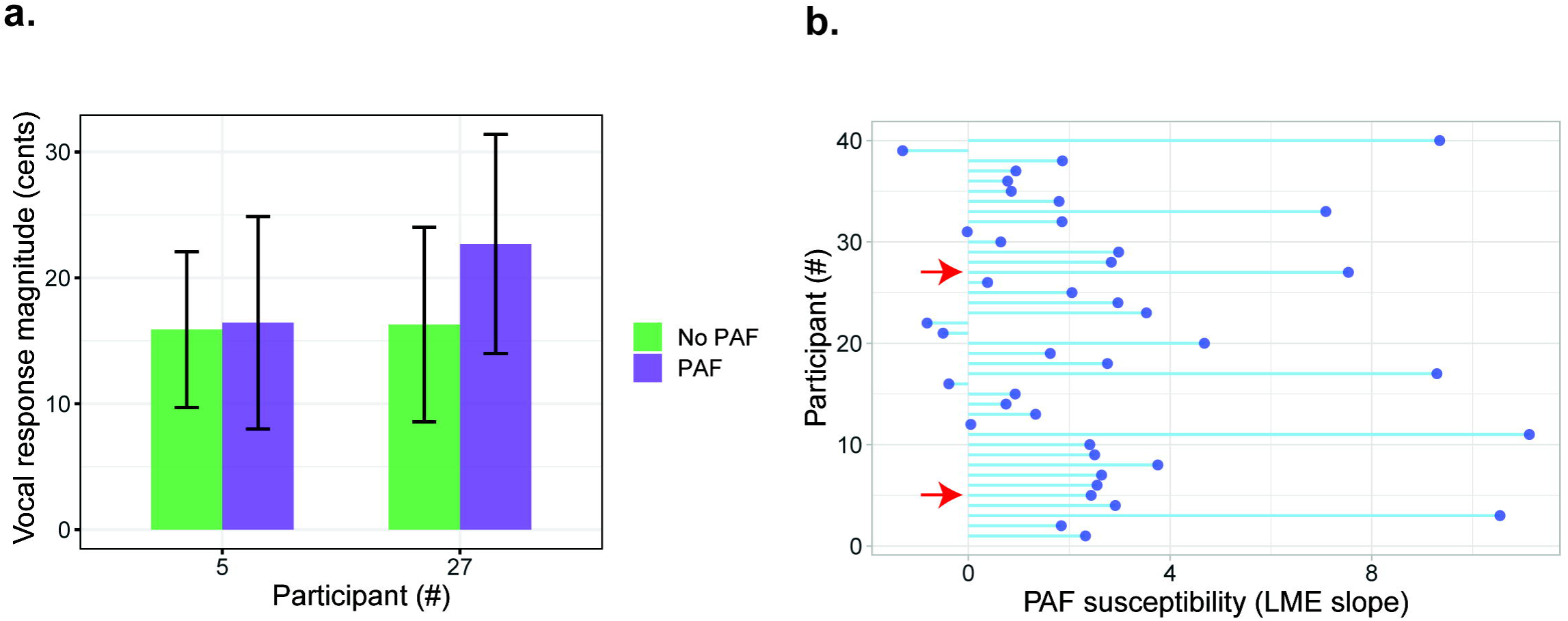
Individual variability in susceptibility to perturbed auditory feedback. **a)** Comparison of vocal response magnitudes during non-perturbed (green bars) and perturbed (purple bars) auditory feedback in two different participants. **b)** Slope values from the LME model which represent the effect of perturbation on vocal response magnitude for different participants. Red arrows indicate the participants that were shown in panel a.

To examine the interindividual variability in susceptibility to perturbed auditory feedback across participants, a susceptibility index was determined for each participant using the slope value of the fixed effect ‘perturbation’ from the fitted LME model. Across participants, there was a considerable variability in susceptibility to perturbation, measured by an increase in vocal response magnitude during perturbation compared to no perturbation (slope values ranging -1.3 to 11.1) (**Figure 7b**).

Finally, to examine whether there is a correlation between the susceptibilities to delayed and perturbed auditory feedback, we compared the susceptibility indices (LME slopes in Figure 3b and 7b) across participants, however there were no significant correlation (Spearman correlation: r = 0.26, p = 0.1). This suggest that an individual can be susceptible to the effects of delayed auditory feedback but not necessarily to the effects of perturbed auditory feedback, or vice versa.

## DISCUSSION

This study highlights that individuals vary in their susceptibility to auditory feedback manipulations. Through the use of both delayed auditory feedback (DAF) and perturbed auditory feedback (PAF) paradigms, we found that susceptibility to temporal manipulations does not necessarily align with susceptibility to acoustic manipulations. These findings imply separate mechanisms at play in speech monitoring for each manipulation type.

Theoretical models of speech production propose that during speech production, motor control regions issue a motor command to the articulators and concurrently transmit an efference copy of this command to the auditory control regions. The efference copy acts as a predictive signal that inhibits the sensory response to one’s own speech. If there is a mismatch between the prediction and the auditory feedback of the produced speech, e.g. when auditory feedback is perturbed, an error signal will be generated to adjust the motor command and correct the speech error (26-28). In this study, we predicted that temporal and spectral auditory feedback perturbations elicit different types of speech errors, which are possibly processed differently. Indeed, a previous meta-analysis study reviewed fMRI investigations that used frequency-altered, delayed, and masked auditory feedback, and found only a small activation overlap in the anterior auditory cortex (29). This observation suggests that distinct neural systems process temporal and spectral aspects of auditory feedback, and are involved in correction of speech errors induced by these different manipulations. Our behavioral results provided further evidence supporting these neuroimaging findings by showing that susceptibility to delayed auditory feedback does not necessarily predict susceptibility to perturbed auditory feedback, such that individuals may be strongly affected by one type of manipulation while remaining largely unaffected by the other.

Our results demonstrated a substantial variability in susceptibility to the effects of auditory feedback manipulations. However, it is not clear what determines the susceptibility of different individuals. Previous studies considered sensory modality preference, suggesting that individuals who rely more heavily on auditory feedback compared to somatosensory feedback during speech production are more susceptible to auditory feedback manipulations (30). To test this idea, a previous study simultaneously altered auditory and somatosensory feedback by applying formant perturbation and mechanical loads to the jaw during speech production. Indeed, there was an inverse relationship between reliance on auditory versus somatosensory feedback, indicating that individuals who compensated more for one perturbation tended to compensate less for the other (20). DAF introduces an asynchrony between the auditory and somatosensory feedback, and compensatory attempts to correct this asynchrony disrupt fluency, resulting in slower speech. In our experiment, we used DAF at 200 milliseconds latency, which is known to result in maximal disruption in healthy individuals, independent of speech rate or the length of utterance (31, 32). Instead of additionally manipulating somatosensory feedback, we tested whether an alternative form of sensory feedback that is synchronous with articulation (i.e. immediate visual feedback) can mitigate the disruptive effects of delayed auditory feedback. We expected that especially less susceptible individuals can adapt strategies to overcome the effects of DAF by disregarding auditory feedback and focusing on visual feedback. However, in contrary to our expectations, immediate visual feedback intensified the disruptive effects of DAF. A prior investigation revealed that introducing delayed visual feedback alongside DAF also intensified the disruptive effects of DAF, suggesting that synchronizing visual feedback with delayed auditory feedback may also not provide a solution (33). These findings illustrate a departure from speech perception, where visual speech cues enhance speech intelligibility in noisy auditory environments (34), as they did not enhance speech fluency under adverse speech production circumstances. This may be because visual feedback is not naturally available during speech production, it is less likely integrated with auditory feedback to aid speech monitoring.

Our results additionally demonstrated that with repeated exposure to auditory feedback, participants exhibited a decrease in their response magnitudes. This phenomenon, termed sensorimotor adaptation, involves updating internal motor plans based on sensory feedback to prevent future discrepancies between predicted and perceived sensory outcomes of motor actions (7, 35). For the DAF experiment, articulation duration of words significantly decreased throughout the course of the experiment with repeated exposure to delayed feedback, confirming findings from previous studies (36). For the PAF experiment, compensatory vocal response magnitudes also decreased throughout the course of the experiment, however this effect was only prominent for 200-cents perturbation condition. Also, 200-cents perturbation elicited fewer opposing responses compared to 100-cent perturbation. Together, these results support the notion that sense of agency might play a role in error correction, whereby compensatory responses occur primarily when auditory feedback is perceived as self-generated. Larger perturbations such as 200 cents are more likely perceived as originating from an external source (37, 38). Therefore, for the PAF experiment, the noted decrease in vocal response magnitudes during 200-cent perturbation is less likely attributed to sensorimotor adaptation. Instead, it could be that participants disregard auditory feedback as self-generated and improve their ability to maintain their voice pitch by merely ignoring it.

Vocal responses within the first 100-150 milliseconds after pitch perturbations are not considered as compensatory and error correction related (39). This is because it takes a longer time for the auditory system to process the vocal output and inform the motor system to make the necessary corrections (40). We therefore focused in the 200 to 500 millisecond period after the perturbation onset to assess compensatory response magnitudes. Closely examining pitch contours for both opposing and following response types revealed that vocal responses to upward and downward perturbations already differed before this time window and even before the perturbation onset. We observed that if the ongoing fluctuation was increasing before perturbation, then vocal response to the perturbation was in the decreasing direction, and vice versa. This pattern was also demonstrated by a previous study that extensively examined vocal pitch fluctuations for opposing and following responses (25). They proposed that continuous vocal pitch fluctuations serve as predictors of the response to perturbation, indicating an interaction between the current state of the speech production system and auditory feedback in determining the type of response. Our findings revealed that a substantial number of responses was following the direction of the perturbation. However, the reasons why participants responded in either direction for the same perturbation type remain unclear. As it is beyond the scope of our study to understand the role of sense of agency or state of the speech production system in determining the response type, we focused exclusively on opposing responses in our analysis to assess a participant’s susceptibility to the auditory feedback perturbation, as these compensatory responses are commonly associated with error correction (7, 35).

In future studies, it will be crucial to investigate the neural mechanisms underlying the correction of different speech errors. Understanding the structural and functional distinctions among different susceptibility profiles and how they translate to disordered populations will be imperative.

## Acknowledgements

We thank Birgit Knudsen, Janniek Wester and Iris Schmits for their support in data collection, and Marteen van den Heuvel for programming the stimulus presentation.

